# Bioplastic production by harnessing cyanobacteria-rich microbiomes for perpetual synthesis

**DOI:** 10.1101/2023.11.06.565755

**Authors:** Beatriz Altamira-Algarra, Artai Lage, Ana Lucía Meléndez, Marc Arnau, Eva Gonzalez-Flo, Joan García

**Author notes:** Corresponding author: Joan Garcia.

## Abstract

Departing from the conventional axenic and heterotrophic cultures, our research ventures into unexplored territory by investigating the potential of photosynthetic microbiomes for polyhydroxybutyrate (PHB) synthesis, a biodegradable polyester that presents a sustainable alternative to conventional plastics. Our investigation focused on a cyanobacteria-enriched microbiome, dominated by *Synechocystis* sp. and *Synechococcus* sp., cultivated in a 3 L photobioreactor under non-sterile conditions, achieving significant PHB production—up to 28% dry cell weight (dcw) over a span of 108 days through alternating cycles of growth and accumulation. Nile Blue staining and Transmission Electron Microscope visualization allowed to successfully confirm the presence of PHB granules within cyanobacteria cells. Furthermore, the overexpression of PHA synthase during the accumulation phase directly correlated with the increased PHB production. Also, gene expression changes revealed glycogen as the primary storage compound, but under prolonged macronutrient stress, there was a shift of the carbon flux towards favoring PHB synthesis. Finally, analysis through proton Nuclear Magnetic Resonance further validated the extracted polymer as PHB. Overall, it was demonstrated for the first time the feasibility of using phototrophic microbiomes to continuous production of PHB in a non-sterile system. This study also offers valuable insights into the metabolic pathways involved.

## 1. Introduction

The increasing concern over environmental pollution and climate crisis has pushed the search for sustainable alternatives to petroleum-based plastics. Bio-based plastics have emerged as a promising solution, offering the potential to mitigate the adverse environmental effects associated with traditional plastics, including the environmental impact of crude oil extraction and the challenges posed by their extremely slow natural degradation [1,2]. Bio-based plastics represent an important alternative to petroleum-based plastics due their organic-based origin [3]. The environmental advantages of bioplastics include reduction in fossil fuel dependency, decreased accumulation of plastic waste, and a diminished carbon footprint [1]. In fact, the demand for biodegradable plastics is rapidly growing, with production set to increase from 2.3 million tons in 2022 to 6.3 million tons by 2027 [4]. Among these alternatives, polyhydroxyalkanoates (PHAs) have gained considerable attention due to their similar mechanical properties to traditional plastics [2]. PHAs are synthesized by various bacteria as a response to inorganic nutrient deprivation, accumulating as intracellular granules. For further insights into bio-based plastics and PHAs, comprehensive reviews are available in the works of [2,5]

Achieving the full potential of bio-based plastics necessitates innovative approaches, and herein lies the promise of cyanobacteria. Cyanobacteria can accumulate polyhydroxybutyrate (PHB, a type of PHA) under nutrient-limited conditions [6–10]. They offer a unique avenue for bioplastic production, harnessing sunlight and carbon dioxide. However, the translation of environmental biotechnologies involving cyanobacteria into practical applications encounters scientific challenges. Despite their recognized potential, particularly within the food industry, the scalability of cyanobacteria cultures and their culture conditions remains a bottleneck [6,11,12]

Notably, the productivity achieved up to now by cyanobacteria wild-type (wt) strains monocultures in autotrophic conditions is not very high, being usually lower than 15 % dry cell weight (dcw) PHB. In a few cases, remarkably high values of up to 20-25 %_dcw_ PHB have been reported in *Synechocystis* sp. and *Synechococcus* sp.[8,13]. To enhance productivity, molecular biology techniques targeting the overexpression of genes implicated in PHB metabolism have been used [14]. However, to make PHB production processes a reality in an industrial context and cost-competitive with the current plastic market, the use of engineered strains seems not to be the most suitable strategy due to the considerable expenses involved in developing and maintaining engineered strains. An alternative procedure involves supplementing cultures with an external organic carbon source, like acetate (Ac). This has led to PHB production of up to 46 %_dcw_ PHB in cyanobacteria monocultures of *Anabaena* sp.[15], 26 %_dcw_ PHB in *Synechoccocus* sp.[8] and 22 %_dcw_ PHB in *Synechocystis* sp.[16]

Nevertheless, monocultures require precise control and sterile conditions, driving up production costs. An option could be the use of microbiomes (or mixed cultures) which in principle could have more stability than single strain cultures when growing in complex media. Microbiomes are potentially more resilient to fluctuations in environmental conditions and less susceptible to contamination with competing microorganisms. Probably, the most well-known microbiome for environmental applications is activated sludge. Studies with microorganism originated from activated sludge under heterotrophic conditions have reported up to 60 %_dcw_ PHB [17,18]. Nevertheless, to the authors’ knowledge, the use of photosynthetic microbiomes enriched with cyanobacteria for PHB production has only been tested in very few studies, including [7,19–21].

A crucial research gap that needs to be addressed in cyanobacteria biotechnology is the maintenance of productive cultures for long-term bioproduct generation. Unfortunately, most experiments to date have been limited to short time, typically lasting only a few weeks, and conducted on a small scale in batch experiments under sterile conditions [10,14,22]. To our knowledge, five cultivations have been reported concerning larger-scale PHB production with cyanobacteria cultures. These include non-sterile tubular photobioreactors inoculated with *Synechocystis* sp. CCALA192 cultivated in 200 L volume [23], a randomly mutated strain of *Synechocystis* sp. PCC6714 cultivated in 40 L volume [24], *Synechococcus leopoliensis* cultivated in 200 L volume [25], wastewater-borne *Synechocystis* sp. in 30 L volume [11], a wild consortium of cyanobacteria was cultivated in a 11.7 m^3^ volume [26]. These cultivations confirm that upscaling cyanobacteria cultivation in closed or semi-closed systems under non-sterile conditions is feasible. However, the PHB production achieved was relatively low, the highest reported was 12.5 %dcw PHB in 75 days [23]. However, considering the commercialization of phototrophic PHB production, it is is imperative to maintain optimal growth conditions, high productivity, resilience to environmental fluctuations, and evaluate economic feasibility to ensure efficient and sustainable large-scale production. In this scenario, adopting sustainable methods, such as developing strategies for recycling nutrients and utilizing waste streams as feedstocks, can reduce operational costs and environmental impact [27,28].

In a previous study, we assessed the viability of augmenting PHB production by enhancing the population of biopolymer-producing organisms via a dual-phase approach involving alternating cell growth on PHB and subsequent biopolymer accumulation induced by Ac addition in a dark environment [21]. Although up to 22

%_dcw_ PHB was obtained after 179 days of operation, the presence of competing green algae resulted in the destabilization of the microbiome and ultimately led to the green algae outcompeting the biopolymer-producing organisms, thus hampering PHB production stability.

In light of the above, in the present study we demonstrate for the first time the capacity of a photosynthetic microbiome enriched in cyanobacteria (and without green algae) to produce PHB over a sufficient extended period to prove perpetual production. To achieve this, we cultivated a microbiome - a diverse microbial culture comprising various cyanobacteria strains and other microorganisms- in a photobioreactor for a total of 108 days, alternating growth/accumulation phases in controlled but non-sterile conditions. Nile Blue A staining and Transmission Electron Microscopy (TEM) were used to visualize intracellular PHB granules inside the cyanobacteria cells. We also analyzed gene expression by quantitative real-time PCR (RT-qPCR) to explore the metabolic pathways involved in PHB synthesis. Finally, polymer characterization was performed by means of Raman Spectroscopy, Fourier Transform Infrared Spectroscopy (FTIR) and proton Nuclear Magnetic Resonance (^1^H-NMR). The integration of diverse analytical techniques offered a comprehensive and multi-dimensional understanding, enhancing the accuracy and depth of research findings.

The results of this study shed light on the long-term capabilities of cyanobacteria microbiomes to generate PHB, which could have significant implications for the bioplastics industry.

## 2. Material and methods

### 2.1. Inoculum and experimental set-up

Two microbiomes isolated in [20], named R3 and UP, were used as the inoculum for 3 L glass cylindrical photobioreactors (PBRs) of 2.5 L working volume (Supplementary Fig. 1). Briefly, microbiome sample R3 was collected from the Besòs River (Sant Adrìa de Besòs, Spain, 41°25’20.2“N 2°13’38.2“E), an intermittent Mediterranean stream that receives high amounts of treated wastewater discharged from the sewage treatment plants in the metropolitan area of Barcelona. UP sample was collected from an urban pond located in Diagonal Mar Park (Barcelona, Spain, 41°24’31.0“N 2°12’49.9“E), which is fed with groundwater. Then samples were cultured in BG-11 medium with low P concentration (0.2 mg⋅L^−1^) to select them over other phototrophs. Cultures grew under 5 klx illumination (approx. 70 µmol m^−2^ s^−1^) with a 15:9 h light:dark photoperiod provided by cool-white LED lights and continuous magnetic agitation. Biomass was scaled up every 15 days using a 1:5 ratio up to 1 L Erlenmeyer flasks. Phylogenetic analysis based on 16S rRNA gene sequences was used to identify the species within the microbiomes [20]. The analysis revealed that R3 exhibited a rich presence of unicellular cyanobacteria, specifically *Synechocystis* sp. and *Synechococcus* sp. (Supplementary Fig. 2A and B), identified as *Synechocystis* sp. PCC6803 and *Synechococcus* sp. PCC 6312, respectively. Sample UP was found to contain *Synechococcus* sp., identified as *Synechococcus* sp. PCC 6312, alongside green algae (Supplementary Fig. 2C and D).

Illumination in reactors was kept at 30 klx (approx. 420 µmol·m^-2^·s^-1^) by 200 W LED floodlight, placed at 15 cm from the reactors. This illumination followed a 15:9-hour light-to-dark cycle during the growth phase. pH levels were continuously monitored using a pH probe (model HI1001, HANNA instruments, Italy) placed inside the reactors. During the growth phase, pH was controlled within a range of 8 ± 0.5 using a pH controller (model HI 8711, HANNA instruments, Italy). When pH levels reached 8.5, a control system activated an electrovalve to inject CO_2_ into the reactors. The pH data were recorded at 5 min intervals using software PC400 (Campbell Scientific). In PHB-accumulation phases (see below), the pH was measured but not controlled in order to avoid IC injection. To ensure darkness during PHB accumulation, the reactors were enclosed in opaque PVC tubes. Reactors were continuously agitated by a magnetic stirrer ensuring a complete mixing and culture temperature was kept at 30-35 °C. Two PBRs were used as duplicates to ensure consistency in the results.

### 2.2. Experimental strategy

Methodology described in [21] based in cycles of alternation of growth/accumulation phases was applied for 108 days (Fig. 1). Briefly, experiment started with a conditioning period consisting on a unique cycle with one growth phase and accumulation phase. The conditioning period was implemented with the aim of promoting optimal conditions for biomass growth, and establishing specific environmental conditions conducive to the subsequent repetitions of the experiment. Specifically, the growth phase started with the inoculation of the PBR with a biomass concentration of 100 mg volatile suspended solids (VSS)·L^-1^. BG-11 with modified concentrations of bicarbonate, as source of IC, N and P (100 mgIC L^-1^, 50 mgN·L^-1^ and 0.1 mgP·L^-1^) was used as media (Table 1). When N was depleted, the starvation phase began. 600 mg Ac·L^-1^ was added at this point and PBRs were enclosed with PVC tubes to avoid light penetration. Note that in this context, we interchangeably used the terms “accumulation” or “starvation” phase to refer to the timeframe during which cells synthesize PHB under nutrient deprivation.

**Figure 1.**
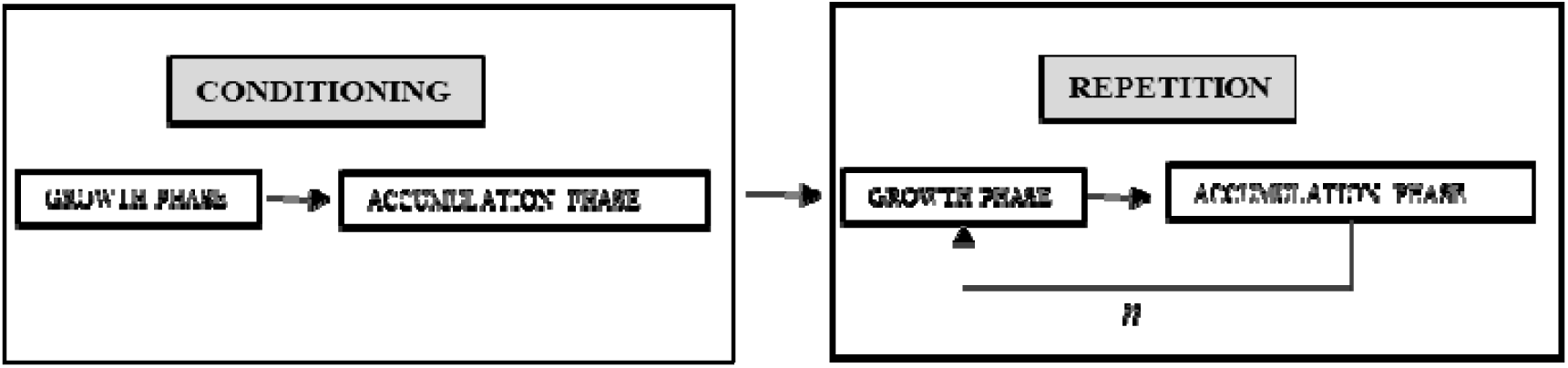
Schematic representation of the methodology applied. *n* is the number of repetitions performed. For microbiome R3, *n* is 4 and for microbiome UP, *n* is 3. Note that here we employed “accumulation” to denote the period when cells synthetize PHB. The word “starvation” is utilized as synonym in the text to refer to this timeframe, as the cells were deprived of nutrients during this period.

**Table 1.** Culture conditions.

| Period | Phase | IC (mg·L<sup>-1</sup>) | N (mg·L<sup>-1</sup>) | P (mg·L<sup>-1</sup>) | Ac (mg·L<sup>-1</sup>) | Lightness<br>(h of light:dark ) |
| --- | --- | --- | --- | --- | --- | --- |
| Conditioning | Growth | 100 | 50 | 0.1 | - | 15:09 |
|  | Starvation | - | - | - | 600 | 0 |
| Repetitions | Growth | - | 25 | 0.1 | - | 15:09 |
|  | Starvation | - | - | - | 600 | 0 |

Following the conditioning period, microbiome R3 underwent a total of five repetitions, while UP underwent three repetitions due to low PHB synthesis attributed to microbiome composition (Supplementary Fig. 2). At the beginning of each repetition, approximately 800 mL to 1,200 mL of culture broth was discarded from the PBRs to purge the system, and replaced with new BG-11 medium with 25 mgN·L^-1^, 0.1 mgP·L^-1^ and without IC, achieving an initial biomass concentration of approximately 400 mgVSS·L^-1^. A daily dose of a solution of KH_2_PO_4_ was conducted to maintain a certain P concentration inside the reactors (aprox. 0.1 mgP·L^-1^). Each growth phase lasted seven days, until N was depleted. After that, the starvation phase started with the addition of acetate to reach 600 mgAc·L^-1^ in the cultures. All the starvation phases went on for 14 days each; except repetition 4 by microbiome R3, which only lasted a week.

### 2.3. Analytical methods

At selected times, 50 mL of mixed liquor were collected. Biomass concentration was determined as VSS according to procedures in [29]. Turbidity was measured with turbidimeter (HI93703, HANNA Instruments). VSS and turbidity were correlated by calibration curve (Supplementary Fig. 3), allowing for a quick estimation of biomass concentration.

To determine the concentration of dissolved chemical species, samples were first filtered through a 0.7 μm pore glass microfiber filter to remove particulates. Nitrate concentration was quantified following method 4500-NO^-^_3_ (B) from Standard Methods [29]. Note that in BG-11 the only source of N is nitrate. The filtered samples were passed through a 0.45 μm pore size filter to determine Ac (acetate) by ion chromatography (CS-1000, Dionex Corporation, USA).

Measurements were conducted in triplicate to ensure robustness and accuracy of the data.

### 2.4. Microscopy

Biomass composition was monitored at the end of each cycle under bright light and fluorescence microscopy (Eclipse E200, Nikon, Japan). Identification and classification of cyanobacteria and green algae were achieved based on their morphological characteristics [30,31]. Cell counting was done in a Neubauer chamber at the end of each starvation phase. Individual cells were counted until reach >400 cells to to ensure a margin of error below 10 % [32].

Intracellular PHB in the biomass was detected through a staining process adapted from [33]. Aliquots of samples from the PBRs were prepared by fixing them to glass slides through heat treatment. These slides were then stained with a 1 % (wt/vol) Nile Blue A solution for 10 min at room temperature. Following the staining procedure, any remaining dye was gently rinsed off with distilled water. Subsequently, an 8 % (vol/vol) acetic acid solution was applied to the slides for one minute at room temperature, after which they were again thoroughly rinsed with distilled water and left to air dry. Finally, the stained samples were examined under fluorescence microscopy at excitation and emission wavelength of 490 nm and 590 nm, respectively.

### 2.5. Transmission Electron Microscope

Sample was taken for Transmission Electron Microscope (TEM) observations at the start of the starvation phase (prior to Ac injection), the fourth day and at the end of the starvation phase in repetition 4, corresponding to days 101, 105 and 108 of the whole experiment. Samples from the reactors (4 mL) were centrifuged (2,000 rpm, 10 min). The supernatant was discarded and the pellet was resuspended in fixative 2% paraformaldehyde and 2.5% glutaraldehyde in 0.1 M PB. Fixation was done at room temperature for 2 h. Then, cells were washed four times in 0.1 M PB. Fixed material was subjected to osmification for 3 h at 4 °C. After that time, samples were washed four times with MilliQ water and stored in 0.1M PB buffer at 4 °C. Samples were dehydrated through a graded ethanol series at 4 °C and gentle agitation (one change of 10 min in 50 % ethanol, two changes of 10 min each in 70, 90 and 96 % ethanol and three changes of 15 min each in 100% ethanol). Samples were embedded using EPON 812 resin kit molds and left 72 h in silicon molds in an oven at 60 °C to polymerize. Ultra-thin sections (100 nm) were cut on a SORVALL MT2-B ultramicrotome with a Diatome 45 ° diamond blade, collected on Formvar-coated 300-mesh coper grids and left to dry 15 h. Finally, samples were stained with UA-zero® and 3 % lead citrate and left to dry 12 h. The sections were examined in a PHILLIPS TECNAI-10 electron microscope operated at 100 kV.

### 2.6. Image processing and analysis

Image analysis was performed using FIJI-ImageJ software. TEM images corresponding to different time points (day 101, 105, and 108) during the accumulation phase of repetition 4 were utilized to measure the size of PHB granules produced in each strain. Prior to measurement, the TEM images were calibrated using a scalebar and the arrow tool to obtain accurate dimensions of the PHB granules.

### 2.7. RNA extraction and quantitative real-time PCR

In repetition 4, samples were collected at the start (prior to Ac injection), the fourth day and at the end of the starvation phase, corresponding to days 101, 105 and 108 of the whole experiment. Samples were collected in triplicates. Methodology was adapted from [34]. Fresh biomass (10 mL) was harvested by centrifugation at 14,000 rpm for 5 min at 4 °C and stored at – 80 °C in an ultra-freezer (Arctiko, Denmark). Frozen cells were homogenized in lysis buffer and TRIzol followed by Bead Beating for cell lysis. Afterward, RNA was isolated using the PureLink RNA Mini Kit (Ambion, Thermo fisher Scientific, Waltham, USA) following the manufacturer’s recommendations. The purified RNA was quantified using a Take3 microvolume plate (Synergy HTX, Agilent, USA). The RNA was reverse transcribed using the RevertAidTM Kit (ThermoFisher Scientific, USA) using 100 ng of total RNA according to manufacturer’s protocol with a combination of the provided oligo (dT) and random hexamer primers (20 μL). The quality and quantity of the cDNA fragments was analyzed using a Take3 microvolume plate (Synergy HTX, Agilent, USA).

Gene expression levels were determined using the qPCR thermocycler Quantstudio 3 (ThermoFisher Scientific, USA). Designed primers described in [34] at 300 nM and the Powerup SYBR master mix (ThermoFisher Scientific, USA) were used. The 16S RNA was selected as the housekeeping gene as the one with lower variability between the different tested conditions. For the results, the mean Ct values were determined using the method from [35] by calculating the average of the triplicate measurements for each condition and gene. The ΔCt was calculated by subtracting the mean Ct value of the housekeeping gene from the mean Ct value of the gene of interest. ΔΔCt is the difference between ΔCt of the day 3, and 7 of accumulation and the ΔCt of day 1 (before adding Ac) as control Ct values. Finally, to calculate the relative fold gene expression level, 2 to the power of negative ΔΔCt according to equation 1:

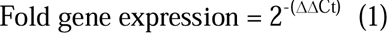

Statistical analysis was performed by one-way ANOVA to evaluate the possible interaction between genes. P-values lower than 5 % were considered statistically significant.

### 2.7. PHB extraction and quantification

PHB analysis was done for samples collected during the starvation phases of both microbiomes. Methodology was adapted from [36]. To begin, 50 mL samples taken from each PBR were centrifuged (4,200 rpm, 7.5 min), frozen at −80 °C overnight in an ultra-freezer (Arctiko, Denmark) and finally freeze-dried for 24 h in a freeze dryer (−110 °C, 0.05 hPa) (Scanvac, Denmark). Approximately 3-3.5 mg of the resulting freeze-dried biomass were combined with 1 mL of methanol solution containing sulfuric acid at 20 % v/v and 1-mL chloroform containing 0.05 % w/w benzoic acid. Samples were heated for 5 h at 100 °C within a dry-heat thermo-block (Selecta, Spain). Following this heating process, the samples were transferred to a cold-water bath for cooling over a period of 30 min. Next, 1 mL of deionized water was added to each tube, which were then vortexed for one minute. The chloroform phase, where PHB had been dissolved, was carefully extracted using a glass pipette and transferred to a chromatography vial equipped with molecular sieves. Analysis of the samples was performed via gas chromatography (GC) (7820A, Agilent Technologies, USA), utilizing a DB-WAX 125-7062 column (Agilent Technologies, USA). Helium served as the gas carrier at a flow rate of 4.5 mL·min^-1^. The injector was set to a split ratio of 5:1 and operated at a temperature of 230 °C, while the flame ionization detector was maintained at a temperature of 300 °C. Quantification of the PHB content was achieved using a standard curve generated from the co-polymer PHB-HV.

### 2.8. PHB characterization

Raman spectra of the samples were acquired using an inVia Qontor confocal Raman microscope (Renishaw) equipped with a Renishaw Centrus 2957T2 detector and a 785 nm laser. All the measurements were performed in mapping mode (64 points) to ensure obtention of representative data. FTIR vibrational studies were recorded on a FTIR Nicolete 6700 spectrometer through a SmartOrbit ATR accessory with Ge crystal and DTGS/CsI detector. Each sample measurement was performed between 4000 – 675 cm^-1^ with a 2 cm^-1^ resolution and spectra processing was carried out using the OMNIC Spectroscopy software. The synthesized polymer and references were analysed through ^1^H-NMR spectroscopy; using a Bruker Avance III-400 spectrometer operating at 400.1 MHz. The chemical shift was calibrated using tetramethylsilane as internal standard and the samples were dissolved in deuterated chloroform (CDCl_3_). Recording of 256 scans was performed for all samples.

### 2.9. Calculations

Total biovolumes (BV) in mm^3^·L^-1^ of each species, including cyanobacteria (*Synechocystis* sp. and *Synechococcus* sp.) and the green algae, were computed using the formula:

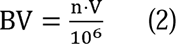

where n represents the count of cells in a sample (cells·L^-1^) and V denotes the average volume of each species (μm^3^). 10^6^ is a factor conversion from µm^3^·mL^-1^ to mm^3^·L^-1^. The cell volumes were estimated using volumetric equations corresponding to the geometric shapes most closely resembling the cells of each species. Specifically, spherical, cylindrical, and ellipsoidal volume equations were utilized for calculating BV of *Synechocystis* sp., *Synechococcus* sp. and green algae, respectively (Supplementary Table 1). Cell dimensions (length and width) were obtained from images of microscope observations (NIS-Element viewer®).

Kinetic coefficients were calculated as follows:

Specific growth rate (d^-1^) was calculated using the general formula

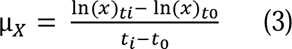

where ln(X)_ti_ and ln(X)_t0_ are the natural logarithms of the biomass concentration (mgVSS·L^-1^) at experimental day (t_i_) and at the beginning of the growth phase (t_0_), respectively. t_i_ values indicate the day when the biomass concentration reaches the stationary phase

Biomass volumetric production rate (mg·L^-1^·d^-1^) was calculated as:

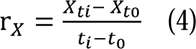

where X_ti_ (mg·L^-1^) and X_t0_ (mg·L^-1^) are the biomass concentration (in mgVSS·L^-1^) at time t_i_ (when biomass reached stationary phase) and at the beginning of the growth phase (t_0_). *i* is the total number of days that the growth phase lasts.

Nitrogen (N) to biomass (X) yield was calculated only during the growth phase by:

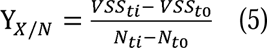

where VSS_ti_ (mg·L^-1^) and VSS_t0_ (mg·L^-1^) denote the biomass concentration at the end (t_i_) and at the start of the phase (t_0_). N_ti_ (mg·L^-1^) and N_t0_ (mg·L^-1^) represent the N concentration (N-NO_3_ ^-^) at the end and at the beginning of each growth phase, respectively.

The specific consumption rate of nitrogen (mgN·mgVSS^-1^·d^-1^) was determined as:

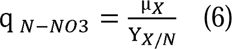

where µ_X_ was obtained as shown in equation 3 and Y_X/N_ in equation 5. PHB volumetric production rate (□_PHB_ (mgPHB·L^-1^·d^-1^)) was obtained by:

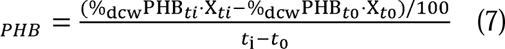

where %_dcw_PHB_ti_ and %_dcw_PHB_t0_ are the percentage of PHB respect biomass quantified at time *i* (end of accumulation phase) and at the beginning of the accumulation phase (t_0_). X_ti_ and X_t0_ are the biomass concentration (in mgVSS·L^-1^) at the beginning (t_0_) and end of the accumulation phase (t_i_).

The PHB yield on acetate (Ac) (Y_PHB/Ac_) was calculated on a chemical oxygen demand (COD)-basis by:

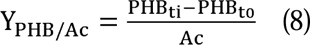

The amount of PHB produced (1.67 gCOD·gPHB^-1^) was obtained by multiplying the %_dcw_ PHB produced per biomass concentration (in mgVSS·L^-1^) at time *i* (end of the accumulation phase) and at the beginning (t_0_) of the accumulation phase. Ac (mg·L^-1^) is the acetate concentration (given 1.07 gCOD·gAc·L^-1^) added (600 mgAc·L^-1^) in the medium at the beginning of the dark phase.

## 3. Results

### 3.1. Consistent growth and PHB accumulation by microbiome R3

The study began by with a first biomass growth phase (conditioning cycle, Fig. 1), wherein two photobioreactors (PBRs) were inoculated with 100 mg volatile suspended solids (VSS)·L^-1^ of microbiome R3 obtained in [20]. A steady-state was reached at the fourth day, when the biomass (as VSS) was approximately 800 mgVSS·L^-1^ (Fig. 2A). The average specific growth rate was 0.52 d^-1^ (Table 2), higher than that obtained with monocultures of *Synechocystis* sp. under similar culture conditions[8,37]. However, it took 18 days for N to be completely depleted (Fig. 2B); likely due to P limitation since it was maintained at a relatively low value (0.1 mgP·L^-1^) to favour cyanobacteria and avoid green algae growth. At this point, the accumulation phase started by adding 600 mgAc·L^-1^ to the medium and enclosing the PBRs with opaque PVC tubes. Starvation phase was maintained 14 days to follow the time course of PHB synthesis by this microbiome. Biomass concentration remained constant during this phase (Fig. 2A). Interestingly, biomass synthetized 11 %_dcw_ PHB during the growth phase, although the conditions were not ideal for biopolymer accumulation due to nutrient presence. Nevertheless, previous studies [22] have reported significant PHB synthase activity, the enzyme involved in biopolymer synthesis, in growing cells of *Synechocystis* sp. PCC6803, the same cyanobacteria strain identified in the microbiome under investigation [20].

**Figure 2.**
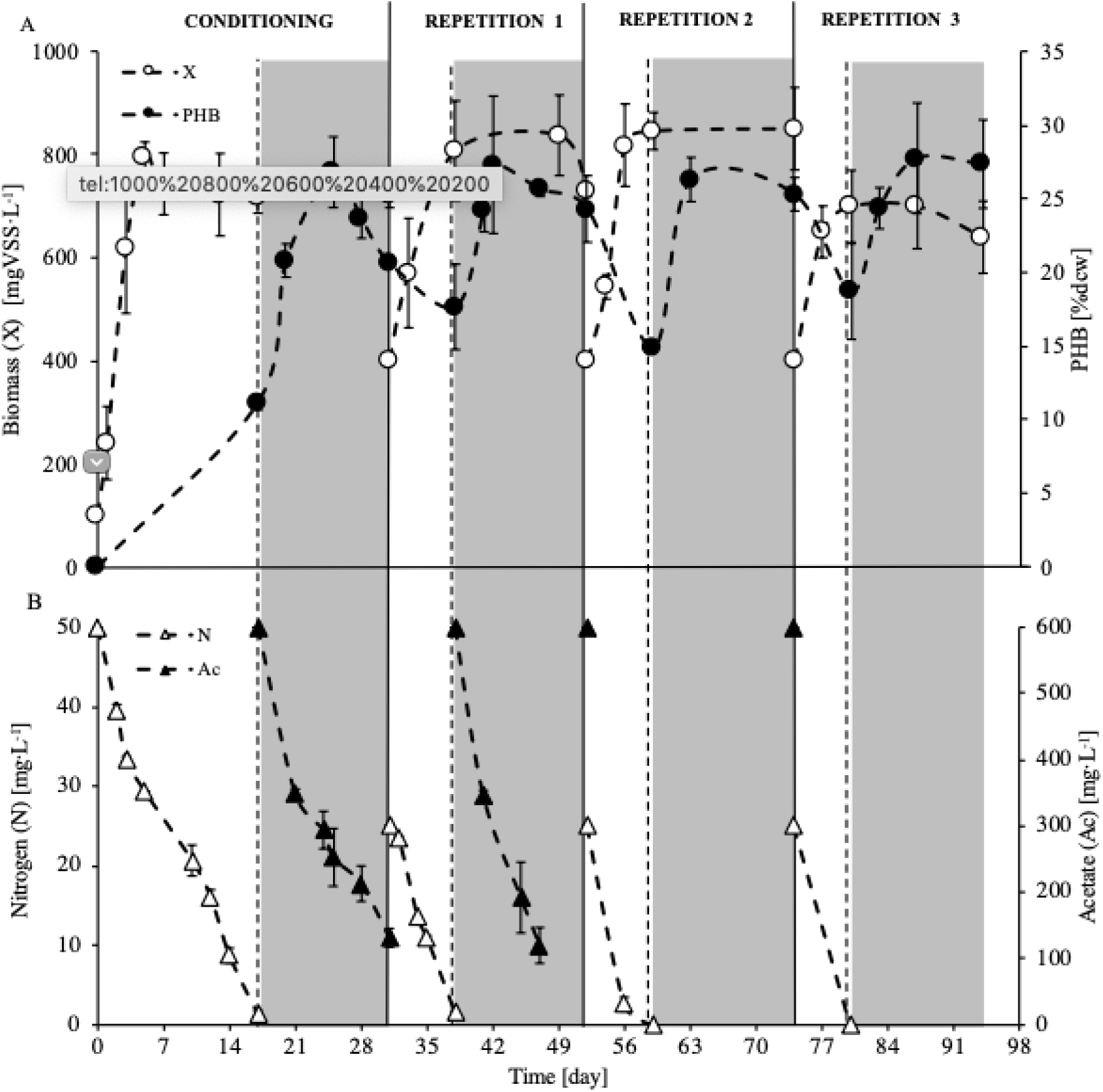
(A) Average biomass (as VSS) and PHB evolution in PBR 1 & 2 of the microbiome R3 for conditioning cycle and repetitions 1-3. Values of biomass were estimated from turbidity measurements. PHB was not measured in growth phase of each cycle. (B) Average nitrogen and acetate evolution through the study for microbiome R3. Ac was not measured in repetitions 2 and 3. Dashed vertical lines indicate the beginning of starvation phase and vertical continuous black lines illustrate end of cycle (conditioning/repetition). Error bars indicate the standard deviation of the replicates, error bars smaller than the symbol are not represented.

**Table 2.** Average of the kinetic and stoichiometric parameters obtained during growth and accumulation phase of each cycle. Values presented in growth phase were measured when biomass reached stationary phase (see Fig. 2). Values in accumulation phase are the average value form both PBRs when the highest PHB content (%dcw) was obtained (at day 8 of accumulation phase in conditioning cycle; day 4 of accumulation phase in repetitions 1 and 2; and day 3 of accumulation phase in repetition 3).

| Growth phase |  |  |  |  |
| --- | --- | --- | --- | --- |
|  | Conditioning | Repetition |  |  |
|  |  | 1 | 2 | 3 |
| VSS (mg·L<sup>-1</sup>) | 802 ± 0.04 | 807 ± 0.09 | 818 ± 0.11 | 650 ± 0.05 |
| μ (d<sup>-1</sup>) | 0.52 ± 0.05 | 0.17 ± 0.04 | 0.18 ± 0.06 | 0.16 ± 0.08 |
| □<sub>X</sub> (mgVSS·L<sup>-1</sup>·d<sup>-1</sup>) | 175.41 ± 10 | 100.02 ± 12 | 104.51 ± 30 | 83.33 ± 15 |
| q<sub>N</sub> (mgN·mgVSS<sup>-1</sup>·d<sup>-1</sup>) | 37.08 ± 5.1 | 6.59 ± 2.3 | 6.39 ± 2.1 | 9.3 ± 1.2 |
| Y<sub>X/N</sub> | 14.03 ± 3.2 | 16.12 ± 2.5 | 16.72 ± 2.3 | 10.15 ± 2.6 |
| Accumulation phase |  |  |  |  |
|  | Conditioning | Repetition |  |  |
|  |  | 1 | 2 | 3 |
| PHB (%dcw) | 27 ± 2 | 27 ± 1 | 26 ± 3 | 28 ± 2 |
| □<sub>PHB</sub> (mgPHB·L<sup>-1</sup>·d<sup>-1</sup>) | 13.55 ± 0.24 | 15.79 ± 0.47 | 17.02 ± 0.65 | 19.74 ± 0.35 |
| Y<sub>PHB/Ac</sub> (g PHBCOD·g AcCOD<sup>-1</sup>) | 0.28 ± 0.02 | 0.16 ± 0.02 | 0.18 ± 0.03 | 0.15 ± 0.03 |

Biopolymer accumulation increased from 11 %_dcw_ to 27 %_dcw_ in seven days, when it reached the maximum. After that, PHB production slowly decreased, since at day 14 (end of the accumulation phase) PHB content was 21 %_dcw_ (Fig. 2A). During this period, biomass consumed 470 mgAc·L^-1^ from the 600 mgAc·L^-1^ added (Fig. 2B).

After 14 days in accumulation phase, a biomass purge was done and replaced with fresh BG-11 medium to start a new cycle (repetition 1) (Fig. 1). Subsequent growth phases (repetition 1, 2 and 3) aimed to select PHB-producers because we assumed that the cyanobacteria will use mostly the stored PHB as carbon source since no substrate (as carbon source) was added to the medium (only the CO_2_ from the injections to maintain pH, Supplementary Fig. 4).

To shorten the growth phase, we used a lower N concentration (25 mg·L^-1^) during the repetitions’ growth phase. This adjustment did not hinder the biomass growth. In fact, biomass reached an average of almost 800 mgVSS·L^-1^ in seven days (Fig. 2A), sufficient for the accumulation step [21]. Biomass exhibited an average growth rate of 0.17 d^-1^ (Table 2) in repetitions 1-3, three times slower than in the first growth performed in the conditioning cycle (µ = 0.52 d^-1^). This difference can be attributed to a lower initial biomass concentration (100 mgVSS·L^-1^ vs. 400 mgVSS·L^-1^), combined with the presence of external IC (as bicarbonate), as well as higher N concentration (50 mg·L^-1^ vs 25 mg·L^-1^) in the conditioning cycle.

Regarding to PHB production, both PBRs followed a similar trend (Fig. 2A). Intracellular PHB increased after Ac supplementation and 14 days in dark, peaking at day 4 of the accumulation phase, when the average was 28 %_dcw_ PHB across repetitions 1, 2 and 3. This corresponds to an average volumetric productivity of approximately 16 mgPHB·L^-1^·d^-1^ (Table 2). Afterward, PHB content decreased but remained relatively constant (around 24 %_dcw_ PHB) for the remainder of the accumulation phase.

pH is useful to track biomass activity. During the growth phase of the conditioning period (initially adding 100 mgIC·L^-1^ as bicarbonate), pH fluctuations were anticipated due to photosynthesis and cell respiration, resulting in daytime rises and nighttime drops in pH (Supplementary Fig. 4). Once the pH reached the setpoint (8.5), CO_2_ was injected in the PBRs to maintain it in the desired range. While 100 mgIC·L^-1^ were present at the start of the conditioning cycle, during repetitions 1-3 bicarbonate was not added. The available IC in the conditioning period enabled more cell grow and; therefore, the increase in pH was faster, resulting in more CO_2_ supplied due to pH control (Fig. 4A). Slower increases in pH during repetitions 1-3 could be attributed to PHB consumption during those growth phases (Fig. 4B). It is difficult to compare pH trends obtained with those from heterotrophic cultures, since often pH is monitored and controlled [8,37,39,40] but pH profiles from the growth phase (referred to as the “famine phase” in heterotrophic cultures) are rarely available.

### 3.2. Presence of green algae overshadowed PHB production

The same methodology described above was applied to two PBRs inoculated with UP, a microbiome rich in cyanobacteria *Synechococcus* sp. and green algae (Supplementary Fig. 2C and D) [20]. In the conditioning period, N (50 mgN·L^-1^) was completely consumed in 25 days, resulting in approximately 550 mgVSS·L^-1^ biomass concentration (Supplementary Fig. 5A) and an average specific growth rate of 0.07 d^-1^ (Supplementary Table 2). This rate was relatively lower compared to that obtained with the microbiome R3, richer in cyanobacteria. Subsequent growth phases (repetitions 1, 2 and 3) were performed without adding bicarbonate to promote growth of PHB-producers. Green algae became noticeable and increased through the experiment, leading to a decrease in the fraction of cyanobacteria in the microbiome population (Supplementary Fig. 6).

Green algae have the ability to accumulate Ac as a carbon storage compound in the form of starch or triacylglycerol under N starvation [41,42] and darkness [43]. Therefore, the abundance of these microorganisms in the PBRs possibly increased because during the accumulation phase (when there was no N or light) they could store the added Ac, competing with cyanobacteria for this compound. They would then use it as carbon source during the subsequent growth phase. Additionally, green algae could grow using the remaining Ac in the PBRs when changing from phase *i* to *i*+1. Around 150 mgAc·L^-1^ remained after 14 days in the accumulation phase of the conditioning cycle (Supplementary Fig. 5B), possibly accounting for the substantial rise in green algae presence during repetition 1. Their proportion increased from 14 % at the end of conditioning cycle to 75 % at the end of repetition 1, and this ratio remained constant for the remainder of the test (Supplementary Fig. 6E).

Regarding to PHB production, unlike microbiome R3 that reached a maximum at day 4, microbiome UP followed a very different trend. PHB increased throughout the 14-day accumulation phase, eventually reaching a maximum value of 7 and 8 %_dcw_ PHB (3 and 4 mgPHB·L^-1^·d^-1^) by the end of the conditioning cycle and repetition 1, respectively (Supplementary Table 2 and Supplementary Fig. 5A). Afterwards, PHB accumulation dropped in repetitions 2 and 3, when only 2 %_dcw_ PHB was detected at the end of these repetitions, representing less than 1 mgPHB·L^-1^·d^-1^ productivity. Differences in PHB production among microbiomes, as well as, its sudden decrease were clearly linked to microbiome composition. Microscope observations showed that after repetition 1, green algae were highly present in microbiome UP (Supplementary Fig. 6), whereas such microorganisms were almost undetected in biomass from R3 (Fig. 3A-B). Such findings suggested that the presence of microalgae overshadowed the potential production of PHB by the microbiome because green algae are non-PHB-producers [21,44,45].

**Figure 3.**
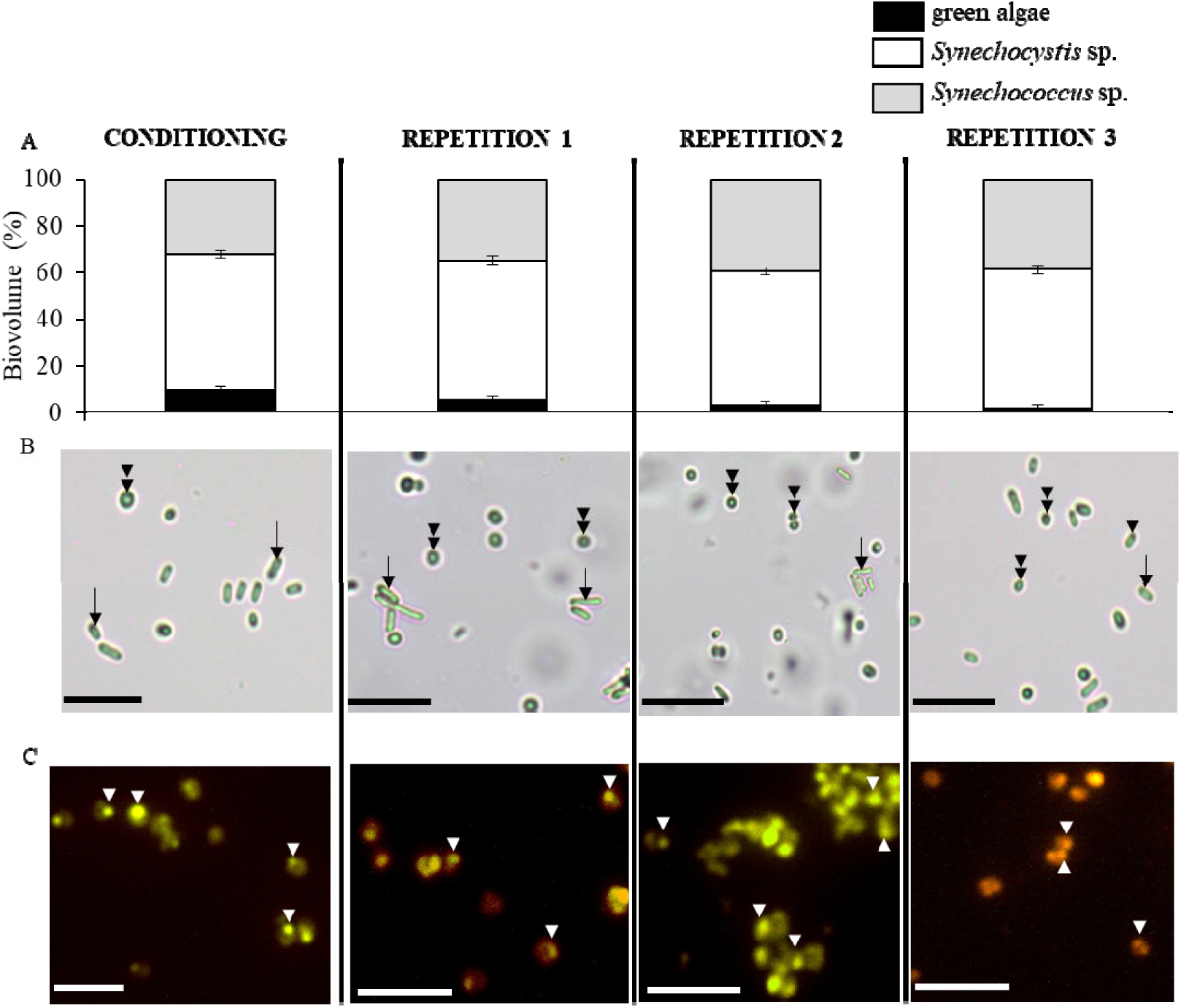
(A) Average biovolume change of different species in microbiome R3. Error bars indicate standard deviation between PBR 1 and PBR 2. (B) Bright light microscope images at 40x of microbiome R3. Double arrowhead points *Synechocystis* sp.; arrow points *Synechoccocus* sp. (C) Fluorescence microscope images at 40x after Nile blue A staining at the end of each cycle. PHB granules were visualized as yellow-orange inclusions after staining. White arrowhead points PHB granules. Each column of images corresponds to the end of the conditioning or end of repetition 1-3, as shown above. Scale bar is 10 µm.

**Figure 4.**
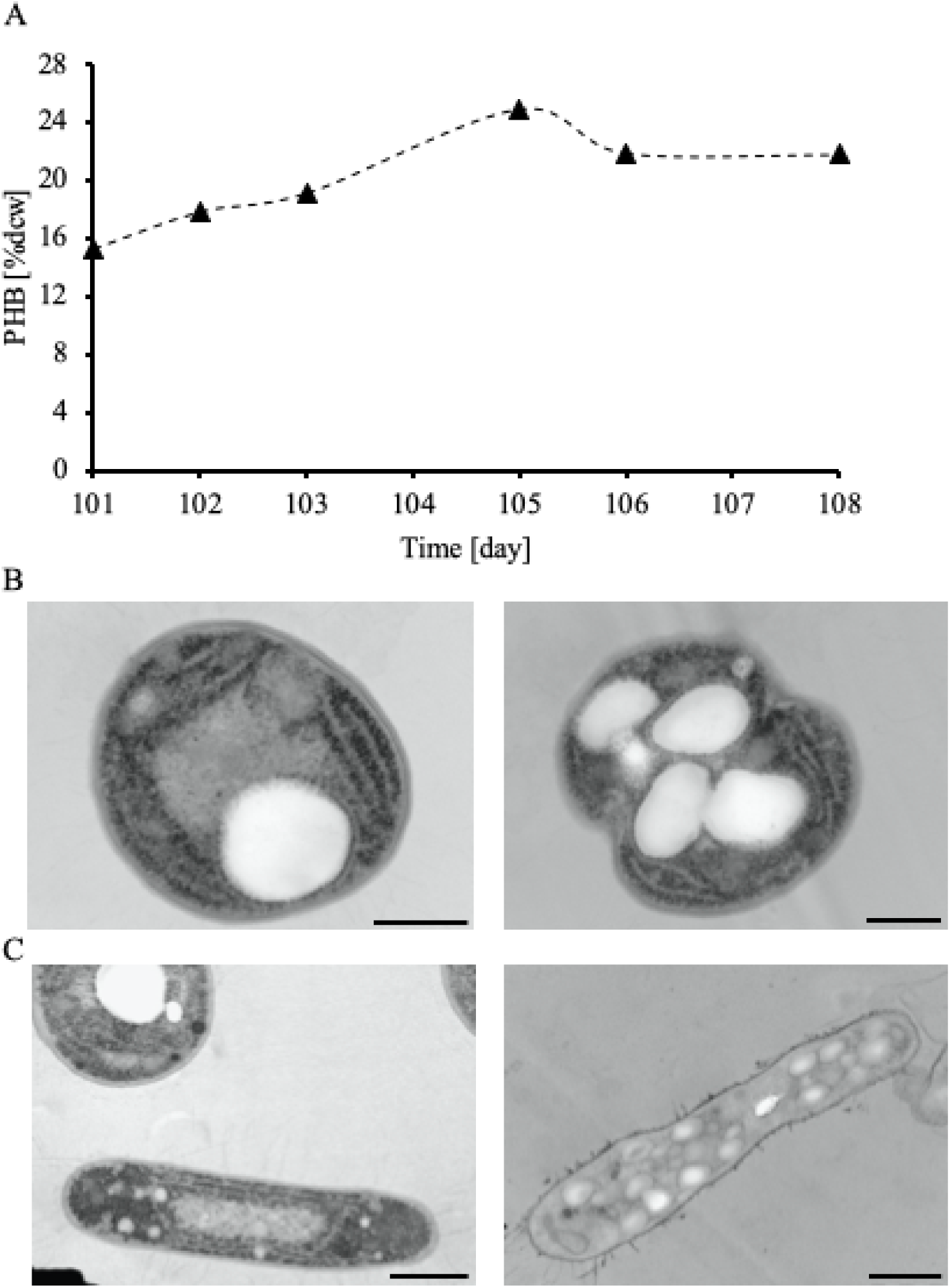
(A) Evolution of PHB production. TEM images of (B) *Synechocystis* sp. And (C) *Synechoccocus* sp. from repetition 4, day 108. In image C left, a *Synechocystis* sp. cell is also visible. PHB granules are visible as white rounded inclusions inside the cell. The heterogeneity of the culture is evident, as there are cells with a high glycogen content (black dots on the thylakoid membrane particularly visible in image C left *Synechococcus* sp. cell); and others where the PHB granules occupy a significant portion of the cellular space. The observable amounts of glycogen in (B) are much larger than those shown in the inoculum, supplementary Figure 7. Scale bar is 500 nm.

### 3.3. Robustness of cyanobacteria microbiome enables high accumulation of PHB

Microscope observations were conducted at the end of each cycle (conditioning, repetitions 1-3) to assess microbiome composition. Outcomes of microbiome R3 showed that the population remained remarkably consistent (Fig. 3A). Notably, an average of 93 ± 2 % of the microbiome comprised two cyanobacteria species, *Synechocystis* sp. and *Synechococcus* sp., indicating a robust and stable microbiome composition in relation to cyanobacteria population. Over the course of the study, both species dominated the culture, although *Synechocystis* sp. was more abundant (60 %) than *Synechococcus* sp. (30 %). In addition, presence of green algae decreased during the operation time; in fact, they were not observed in the microscope observations performed (Fig. 3B).

To verify accumulation of PHB by cyanobacteria and no other microorganisms, Nile blue A staining was performed in samples from the end of each cycle (conditioning and repetitions 1-3). PHB was detected using a fluorescence microscope. The positive staining with Nile blue A clearly demonstrated that cyanobacteria were involved in PHB accumulation (Fig. 3C).

### 3.4. High intracellular PHB content revealed by TEM

A subsequent cycle (repetition 4) of seven days of growth and seven days under starvation was done with one PBR to obtain images of the intracellular PHB-granules. Samples were collected at three time points: at the start (prior to Ac injection), the fourth day (when maximum biopolymer production occurred) and at end of the starvation phase. These time points corresponded to days 101, 105 and 108 of the entire experiment, and are referred by those numbers in this section.

Biomass used as inoculum was also examined by TEM to observe and compare morphological changes in response to the continuous growth/starvation cycles performed. Inoculum cells, grown in BG-11 medium with 0.5 mgP·L^-1^, displayed a typical cyanobacteria cell organization (Supplementary Fig. 7A), with the thylakoid membranes occupying most of the cytoplasm volume. Small electron-dense glycogen inclusions between the thylakoid layers could be seen. Some cells also presented slightly electron-dense inclusions located close to the thylakoid membranes. These inclusions were not PHB nor polyphosphate granules since both have different morphology and electron density after staining [46,47]. Additionally, PHB quantification by gas chromatography (GC) revealed that inoculum had no intracellular PHB. These spherical granules could be presumably carboxysomes and/or lipid bodies.

TEM images of samples taken during the accumulation phase revealed distinct electron-transparent inclusion bodies (“white”) with a transparent appearance, located near the cell periphery, around the thylakoid membranes (Fig. 4B-C and Supplementary Fig. 7). These were attributed to PHB-granules. At the phase’s onset (before adding Ac, day 101), cells already contained PHB granules because they had experienced 4 cycles of growth/starvation (conditioning + repetitions 1-3), and not all PHB was consumed during the growth phases. In fact, PHB quantifications showed that 15 %_dcw_ still remained in the biomass (Fig. 4A, day 101). Notably, *Synechocystis* sp. cells exhibited in general no more than 3 PHB granules at day 101, increasing on day 105 and 108 (Supplementary Fig. 7). Indeed, the highest PHB accumulation was observed on day 105, four days after Ac supplementation (24 %_dcw_ PHB), with *Synechocystis* sp. presenting a maximum of 6 granules per cell (Fig. 4B), while *Synechococcus* sp. cells had up to 15 granules or more (Fig. 4C). On day 108, after 7 days in starvation, no differences were detected in the size and number of PHB-granules per cell with sample from day 105 (Supplementary Fig. 7B). Remarkably, relatively similar PHB content was also detected on both days (24 %_dcw_ PHB on day 105 and 22 %_dcw_ PHB on day 108) (Fig. 4A). PHB-granules had spherical to oval shape in both cyanobacteria species; but the granules were larger in *Synechocystis* sp. compared to *Synechococcus* sp., with average diameters of 672 ± 83 nm and 217 ± 19 nm, respectively.

### 3.5. Expression of key genes involved in PHB metabolism

RT-qPCR was performed to analyze the expression of specific genes encoding key enzymes related to the metabolism of PHB. Samples were analysed in repetition 4 at the same time points in which TEM images were taken (previous section). These are the start of the starvation phase (before Ac injection), the fourth day (when maximum biopolymer production occurred), and the end of the starvation phase, corresponding to days 101, 105, and 108 of the entire experiment. Additionally, enzymes involved in glycogen metabolism were also analysed since PHB can be synthesized from intracellular glycogen pools[7,48]. Both pathways, as well as the TCA cycle, use Acetyl-CoA, which can be synthesized from Ac, as a primary precursor. Results from day 101 served as reference to compare with outcomes from day 105 and 108 (Supplementary Fig. 8). Note that RT-qPCR targeted *Synechocystis* sp. genes, given their high conservation among species [49], and to their dominance in the culture as evidenced by microscope observations (Fig. 3A-B).

On day 105 (fourth day of the accumulation), the overexpression of genes related to glycogen synthesis (*glgA*, codifying for glycogen synthase), the TCA cycle (*gltA*, codifying for citrate synthase) and PHB synthesis (*phaC*, codifying for polyhydroxyalkanoate synthase) was revealed (Fig. 5A). On day 108 (seventh day of the accumulation), genes *phaC* and *glgp1* (codifying for glycogen phosphorylase, involved in glycogen catabolism) were overexpressed (Fig. 5B). The consistent overexpression of *phaC* on the fourth and seventh days of the starvation period (days 105 and 108 respectively) aligns with the observed stable PHB content (24 %_dcw_ PHB and 22 %_dcw_ PHB, on days 105 and 108 respectively).

**Figure 5.**
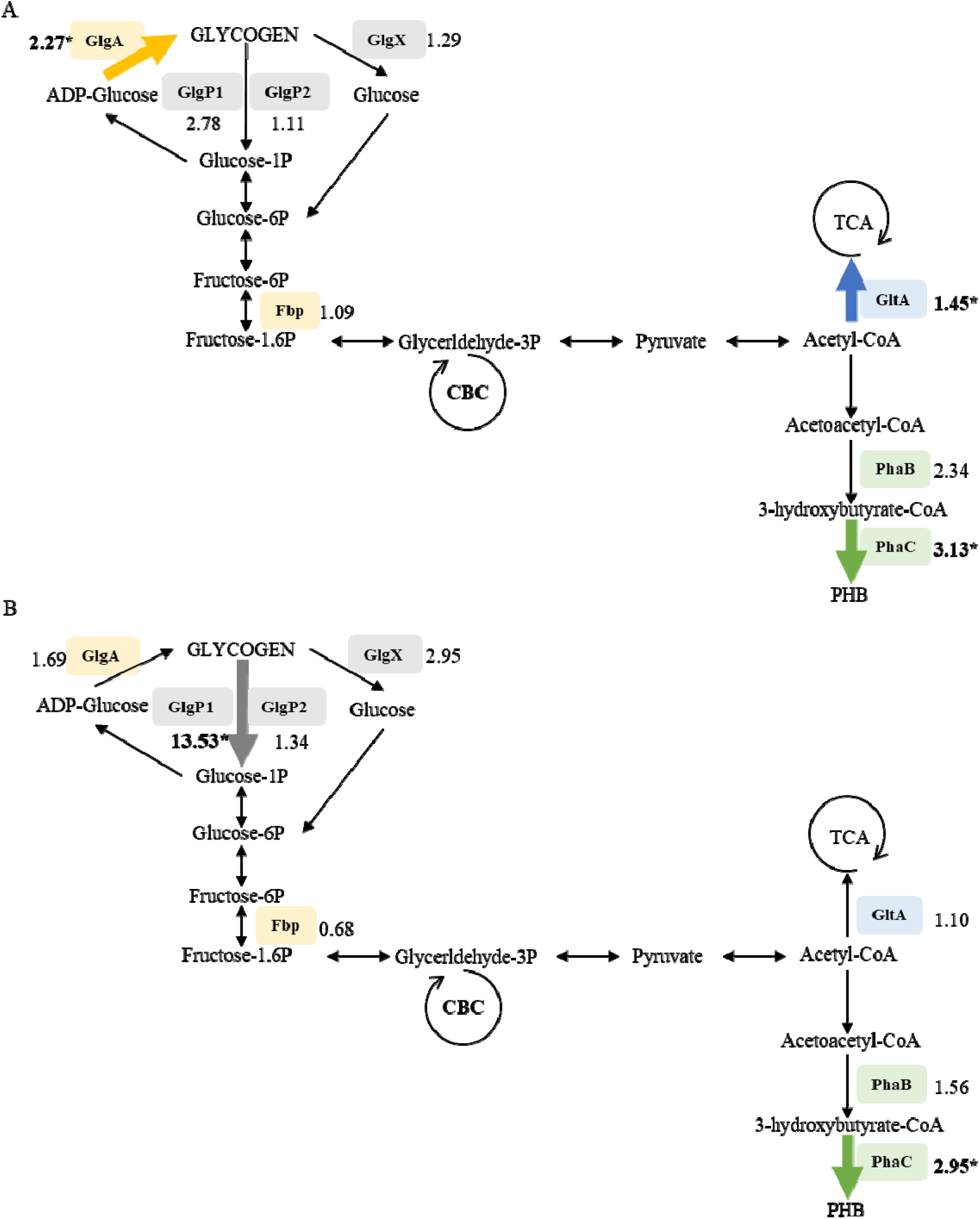
Schematic representation of biosynthetic pathways for PHB and glycogen production in cyanobacteria. (A) corresponds to results from day 105 and (B) day 108. Enzymes codified by the studied genes are shown in squares. Numbers next to enzymes names represent the fold gene expression. * denotes genes with statistically significant overexpression (p-value < 0.05). Yellow shows key enzymes involved in the synthesis of glycogen (Fbp, GlgA); grey, to the glycogen catabolism (GlgP1, GlgP2, GlgX); green, to the synthesis of PHB (PhaB, PhaC); and blue, to the introduction of acetyl-CoA into the tricarboxylic acid (TCA) cycle (GltA). Abbreviations: Fbp: fructose-bisphosphatase 1 GlgA: glycogen synthase, GlgP1 and glgP2: glycogen phosphorylase, GlgX: glycogen debranching enzyme, GltA: citrate synthase, PhaB: Acetyl-CoA reductase, PhaC: poly(3-hydroxyalkanoate) synthase, TCA: tricarboxylic acid cycle, CBC: Calvin-Benson cycle.

### 3.6. PHB characterization

The cyanobacteria-generated biopolymer was assessed with spectroscopic techniques to make a characterization of the composition of the polymer. As reported in Fig. 6A, main Raman active modes for a reference sample of PHB (PHB-R) were observed at 840 (v_1_), 1060 (v_2_), 1300 – 1500 (v_3_), 1725 (v_4_) and 2800 – 3100 cm^-1^ (v_5_) and attributed to C–COO, C–CH_3_ stretching, CH_2_/CH_3_ bending (symmetric and antisymmetric), C=O stretching and different C–H stretching of methyl groups, respectively [50]. Raman spectra comparison between PHB-R and the PHB biogenerated (PHB-B) showed no differences, except from a broad shoulder at 2876 cm^-1^ attributed to impurities acquired during the extraction process (Diamond mark, Fig. 6A). FTIR outcomes (Fig. 6B) corroborated the results obtained by Raman through the observation of the main vibrational modes C–CH_3_ stretching, CH_2_ wagging and C=O stretching (1057, 1281 and 1724 cm^-1^, respectively) reported for PHB [50,51]. A broad band at 3000 – 4000 cm^-1^ caused by water presence was detected for the PHB-B sample leading to a poor signal-to-noise ratio at the same region.

**Figure 6.**
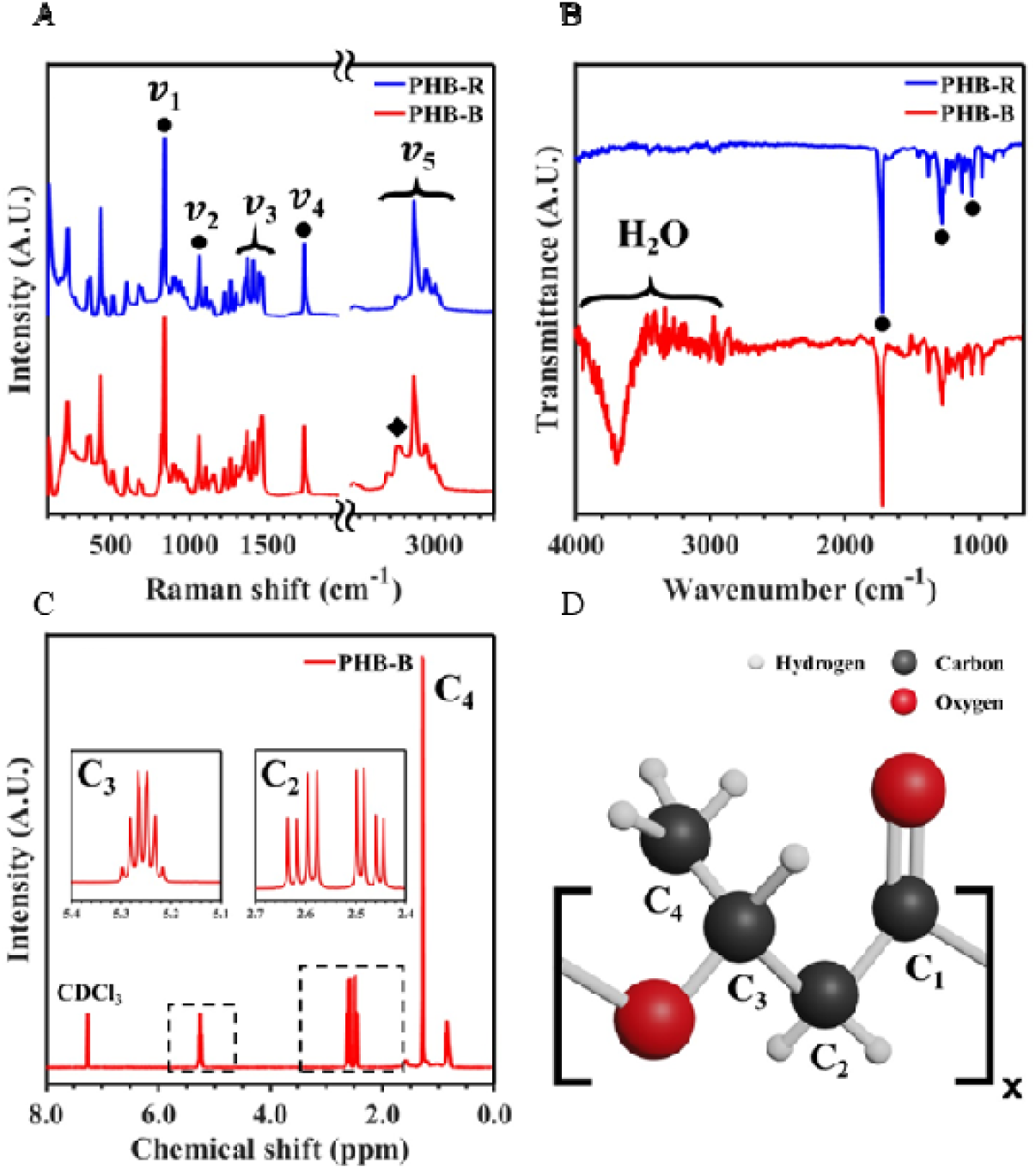
(A) Raman spectra for the PHB-R and PHB-B samples where the main Raman active modes are marked with circles. The diamond highlights the shoulder attributed to impurities during the extraction process. (B) FTIR spectra for the samples PHB-R and PHB-B, where the region affected by the water and the vibrational modes (marked with circles) can be observed. (C) ^1^H-NMR spectra for the PHB-B sample with insets of the relevant peaks and carbon assignation related to the monomer carbons. (D) Schematic drawing of the PHB monomer with carbon numeration for NMR spectra interpretation.

Nonetheless, due to significant similarities in Raman and FTIR spectra between PHB and other PHAs, such as poly(3-hydroxybutyrate-co-3-hydroxyvalerate) (PHBHV, Supplementary Fig. 9A-B), ^1^H-NMR analysis was deemed necessary to further confirm the sole production of PHB. Careful inspection of the PHB-B NMR spectra (Fig. 6C) enabled the peak assignation of the carbons depicted in the PHB monomer (Fig. 6D).

## 4. Discussion

Research on PHA production by bacteria is extensive, but studies involving mixed cultures with cyanobacteria are relatively limited. While results on PHB synthesis by cyanobacteria pure cultures have demonstrated their potential as biopolymer producers [7–10,13,15,16], current production yields may not yet meet the demands of a market predominantly dominated by petroleum-based plastics. Therefore, efforts to boost their productivity should be pursued.

Here, we demonstrate the feasibility of continuous PHB synthesis using a microbiome rich in cyanobacteria. This microbial culture encompassed various cyanobacteria strains and microorganisms, with cyanobacteria driving the PHB production process. This is achieved through the implementation of repetitive biomass growth and PHB accumulation phases. Several key factors contribute to the success of our approach. Firstly, the composition of the microbiome proved crucial for maintaining PHB synthesis over time, with cyanobacteria, the primary PHB producers, requiring dominance in the culture. This strategic control resulted in notable 25-28 %_dcw_ PHB, ranking among the highest values recorded by the cyanobacteria strains present in the studied community (*Synechocystis* sp. PCC 6803 and *Synechococcus* sp. PCC 6312, Table 3). Although PHB synthesis by *Synechocystis* sp. PCC 6803 has been thoroughly investigated, there is a dearth of literature available on the performance of *Synechococcus* sp. PCC 6312, nor the use of both strains in a mixed culture. Previous studies reporting PHB synthesis usually operated with monocultures under sterile conditions and in very small volumes, rarely exceeding 150 mL. In the only report of *Synechocystis* sp. PCC 6803 tested in higher volumes, an engineered strain (ΔSphU) was cultivated with shrimp wastewater in a 15 L PBR [52]. Despite reporting high intracellular PHB content, engineered strains are not optimal for the scale-up of the process in an environmental biotechnology perspective, since production costs will increase due to the requirement of sterile conditions or synthetic substrates.

**Table 3.** Comparison of PHB production in cyanobacteria *Synechocystis* sp. PCC6803.

| Genotype | Working volume (L) | Culture conditions | Accumulation time (days) | PHB fraction (%dcw) | Reference |
| --- | --- | --- | --- | --- | --- |
| WT | 0.1 | N-, P- & Ac+ | 21 | 33 | [80] |
| WT (microbiome) | 3 | N-, P- & Ac+ | 14* | 27 | This study |
| WT | 0.05 | P- & Ac+ | 14 | 26 | [13] |
| WT | 0.05 | N-, P- & Ac+ | 20 | 20 | [81] |
| WT | 0.15 | N-, P- & Glc+ | 12 | 13 | [10] |
| ΔPirC and OE PhaAB (*Cupriavidus necator*) | 0.05 | N-, P- & Ac+ | 20 | 81 | [81] |
| ΔPirC and OE PhaAB (*Cupriavidus necator*) | 0.05 | N- & P- | 20 | 63 | [81] |
| OE PhaAB (native) | 0.05 | N- & Ac+ | 9 | 35 | [14] |
| ΔSphU | 15 | Shrimp wastewater | 11 | 33 | [52] |
| ΔSphU | 0.05 | N+ | 14 | 15 | [82] |
| OE Xfpk | 0.08 | N- & P- | 30 | 12 | [83] |
| OE PhaAB (*Cupriavidus necator*) | 0.05 | N- & Ac+ | 8 | 11 | [22] |

WT: wild-type; OE: overexpression; Δ: deletion; PirC: PII-interacting regulator; PhaA: beta-ketothiolase; PhaB: acetoacetyl-CoA reductase; SphU: phosphate regulator; Xfpk: phosphoketolase; N: Nitrogen; P: Phosphorus; Ac: Acetate; Glc: Glucose; -: defficiency; +: supplementation. \*Note that all references are batch experiments, while in this study, three iterated accumulation phases have been performed with the same culture biomass, representing a total of 108 days of reactor operation.

Our study represents a notable advancement by utilizing a microbiome to synthesize PHB within a 3 L PBR, a departure from previous studies conducted at much smaller scales (Table 3). Our noteworthy accomplishment of sustaining PHB production over a 108-day period highlights the microbiome’s capacity for prolonged, sustainable bioproduction, suggesting promising commercial potential. However, it is crucial to distinguish between the duration of our study and the actual time required for commercial-scale PHB production. Factors like reactor capacity, purification methods, and logistical considerations will significantly influence the production cycle’s real-world timeline. Nonetheless, our study’s extended duration of consistent PHB production underscores the resilience of the microbial culture in accumulating the desired bioproduct. This extended duration of steady production attests to the robustness of the culture in accumulating the desired bioproduct. Moreover, it is noteworthy that cyanobacteria microbiomes present a game-changing alternative to heterotrophic cultures by harnessing inorganic carbon (CO_2_) and sunlight for growth, eliminating the need for energy-intensive aeration, which accounts for 50 % of the process’s energy demand [23,53,54]. In our methodology, PBRs have worked under non-sterile conditions in semi-continuous mode, requiring minimal manipulations, such as Ac addition at the start of each starvation phase and culture purge followed by new medium replacement every 21 days (in fact the purge is the resulting product of our process). By cultivating cyanobacterial microbiomes in industrial bioresources without requiring sterile conditions, operational costs can decrease by up to 40 % [55,56].

We also showed that to ensure optimal PHB production, it is imperative to prevent presence of non-PHB producers, like green algae, in the initial inoculum. Despite the low P concentration used to prevent their proliferation during growth phases, it became evident that the stored carbon in the form of starch or triacylglycerol and/or the residual Ac significantly contributed to their growth (Supplementary Fig. 6). This indicated that P limitation alone was insufficient to hinder the growth of green algae, and other approaches should be included, like manipulating light color (wavelength), to promote cyanobacteria dominance [57,58]. In our previous study, up to 22 %_dcw_ PHB was obtained by a microbiome rich in cyanobacteria; nevertheless, production was marked by notable fluctuations due to the presence of green algae [21]. This underscores the critical role of culture composition in achieving stable and reliable PHB production.

Secondly, we provide compelling evidence supporting the active accumulation of the biopolymer by cyanobacteria, as confirmed through both Nile blue A staining and TEM images (Fig. 3 and 4). In addition, Nile blue A staining could be used as a rapid and effective methodology to asses PHB synthesis in microbial cultures by correlating the fluorescence intensities of Nile blue A and PHA concentrations, aligning with previous reports in sludge from wastewater treatment plants [40,59–61] or the cyanobacteria *Nostoc* sp.[9].

PHB granules were detected in cyanobacteria cells as white inclusion bodies in TEM images. Interestingly, TEM images depicting cyanobacteria cultures with intracellular PHB are not commonly reported. In fact, only a few studies have described the granule size, number, or intracellular content in these microorganisms (Table 4). Most published works focus on morphological changes between WT and mutant cells, where PHB production was not the main objective [47,62,63]. However, based on the available TEM images, we can conclude that the *Synechocystis* sp. from our studied mixed culture exhibited one of the highest intracellular biopolymer contents, as well as the greatest number of granules. Much limited information is found regarding PHB production by *Synechoccocus* sp. From TEM images by [64] and the current study (Fig. 4), *Synechoccocus* sp. presented a higher number of granules, but smaller in size, compared to *Synechocystis* sp. (Table 4). The significant variability observed in both the size and quantity of biopolymer granules across different species and even within cells of the same species has contributed to a heterogeneous biopolymer content in the culture (Supplementary Fig. 7B). Notably, under conditions favourable for PHB synthesis, certain cells failed to accumulate the polymer, instead presenting glycogen as carbon storage compound (Fig. 4B-C). Conversely, in some cells, PHB granules occupied a significant portion of the cellular space. This diversity may be attributed to the stochastic regulation of PHB synthesis [30].

**Table 4.**
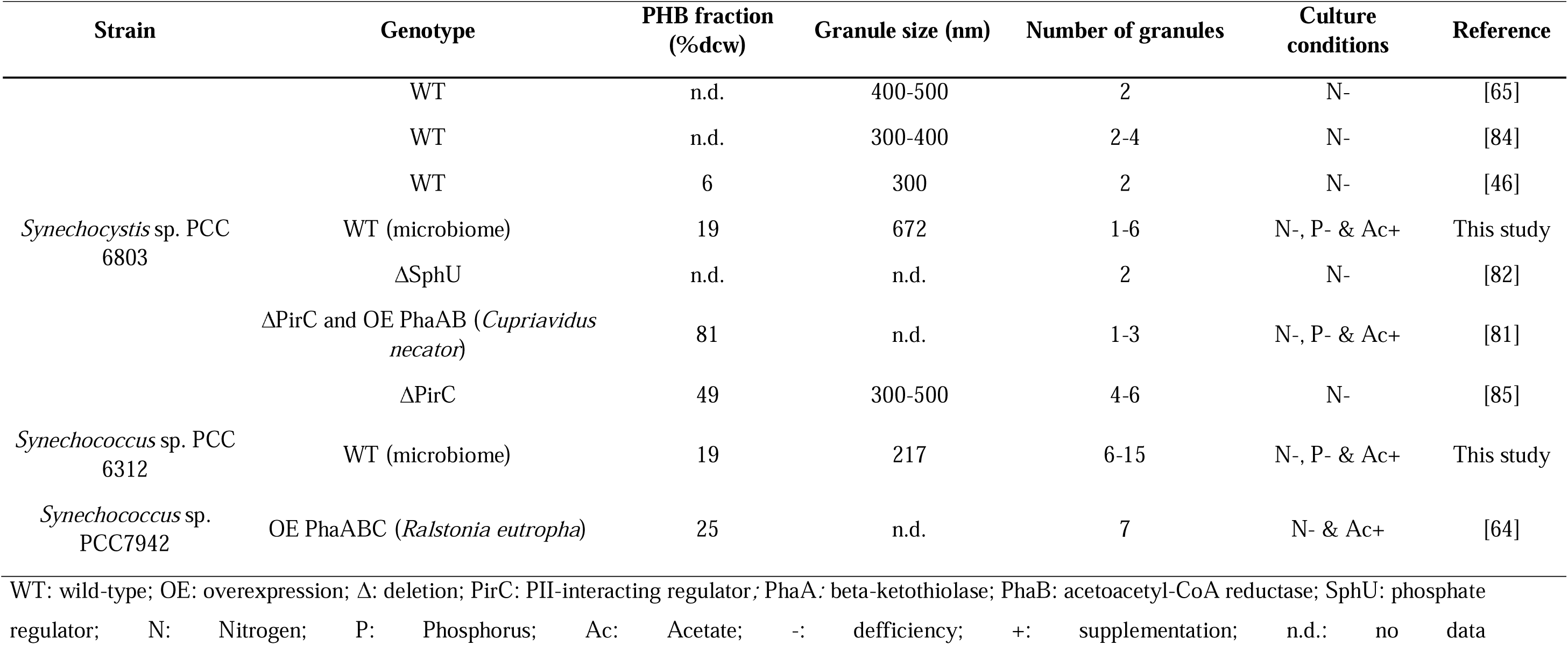
Comparison of published data related to PHB granules in cyanobacteria *Synechocystis* sp and *Synechoccocus* sp.

Thirdly, the direct correlation between the overexpression of the *phaC* gene and increased PHB production during the accumulation phase underscores PhaC’s role as the key enzyme in PHB synthesis. For instance, previous studies in *Synechocystis* sp. have demonstrated that the absence of *phaC* (Δ*phaC*) hindered PHB production when acetate was introduced to the medium[65]. In our investigation, a clear relationship emerged between the increase in PHB production (Fig. 5A) and the overexpression of the *phaC* gene, observed from day 101 to day 105 and 108 (Fig. 5). This temporal alignment, particularly the similar overexpression of *phaC* on the fourth and seventh day of starvation (days 105 and 108, respectively), correlates with a sustained PHB content (24 %_dcw_ PHB in day 105 and 22 %_dcw_ PHB in day 108). Furthermore, the presence of a comparable number of cells containing PHB granules on day 105 and 108 (Supplementary Fig. 7) suggested that polymer synthesis from acetate was a relatively fast process reaching its maximum in four days and remaining constant thereafter.

In addition to PHB, cyanobacteria also store glycogen as carbon storage compound [7,65]. Our findings indicated ongoing glycogen synthesis on day 105, supported by the overexpression of *glgA* (Fig. 5A and Supplementary Fig. 8), which subsequently decreases by day 108 (Fig. 5B and Supplementary Fig. 8). This trend is further supported by the TEM images, wherein certain cells exhibit small electron-dense glycogen inclusions between the thylakoid layers (Fig. 4 and Supplementary Fig.7). The dynamic gene expression suggested that glycogen served as the initial storage compound, synthesized in response to short-term macronutrient stress conditions, such as nitrogen depletion, as reported by other authors [34,65–67]. Nevertheless, the elevated expression of *gltA* on day 105 (Fig. 5A and Supplementary Fig. 8) indicated that a portion of Acetyl-CoA was channeled into the TCA cycle instead of being used for PHB production. Interestingly, this gene was not overexpressed on day 108, implying a decrease in carbon flux to TCA, possibly favoring PHB synthesis.

By day 108, cells had accumulated 22% _dcw_ PHB, corroborated by biopolymer extraction results, and the overexpression of *phaC* (Fig. 4A and Fig. 5B). Notably, no overexpression of genes related to glycogen synthesis was detected at this point, further supporting rapid synthesis of glycogen due to nutrient starvation and subsequent PHB accumulation. Additionally, active glycogen degradation, indicated by the overexpression of the *glgp1* gene (Fig 5B and Supplementary Fig. 8), suggested the ongoing conversion of stored glycogen to PHB, prompted by persistent nitrogen starvation. This metabolic shift explained the sustained intracellular PHB content after 7 days in starvation, supported by the overexpression of *phaC* and the absence of PHB decrease. These findings align with previous studies reporting the conversion of glycogen to PHB in cyanobacteria [34,48,68]. Importantly, the degradation of glycogen and its transformation into PHB during prolonged N-starvation serve to mitigate the potential osmotic impacts of excessive intracellular metabolites accumulation and generate ATP to sustain basic cellular functions [69,70]. This elucidated the metabolic dynamics underlying PHB synthesis.

Finally, to bridge the gap between laboratory-scale production and industrial applicability, the utilization of spectroscopic techniques becomes indispensable for comprehensive material analysis and characterization. With this aim, we conducted Raman, FTIR and ^1^H-NMR analysis to characterize the synthetized biopolymer. Results clearly showed that the biopolymer accumulated by *Synechocystis* sp. and *Synechococcus* sp., the cyanobacteria present in the studied microbiome, under acetate supplementation was PHB. Notably, the absence of peaks with possible attribution to other polymers, coupled with the additional ^1^H-NMR measurements conducted on commercial PHB and on copolymer poly(3-hydroxybutyrate-co-3-hydroxyvalerate) PHBV (Supplementary Fig. 9C-D) provided conclusive evidence for the characterization of the biopolymer as PHB. Note that cyanobacteria can synthetize the copolymer PHBV by the addition of other carbon sources to the medium, such as valerate or propionate [71]. Further comprehensive analysis could be undertaken to investigate the mechanical properties of the biopolymer and explore new potential applications. Examples of studies regarding to this topic include utilizing PHB nanoparticles for drug delivery [72] or exploring combinations with other materials to extend its applications, such as PHB blending with more ductile biopolymers, like poly(lactic acid), starch, cellulose and poly(caprolactone) [73–77], with maleic anhydride [78], or phenol mixtures from winery residues [79].

## 5. Conclusion

In this study, a phototrophic microbiome was harnessed to produce PHB over 108 days, employing alternating growth and accumulation phases. Results demonstrated that a microbiome rich in cyanobacteria achieved a remarkable accumulation of 25-28 %_dcw_ PHB. This achievement stands as one of the highest reported contents in wild-type cyanobacteria over an extended timeframe. Notably, PHB production decreased when green algae were dominant in the microbiome. Additionally, positive Nile Blue A staining and TEM revealed the intracellular location of PHB granules within cyanobacteria cells.

Furthermore, gene expression data offered insights into the metabolic pathways and regulatory mechanisms involved. The overexpression of gene *phaC* exhibited a direct correlation with the increased PHB production during the accumulation phase. The upregulation of genes associated with glycogen metabolism (*glgA* and *glgp1*) pointed to the significant interplay between these storage polymers as essential carbon sources.

Understanding mechanical properties of biopolymers obtained through biological processes is crucial for envisioning broader applications. In our study, as a proof-of-concept, spectroscopic analysis (Raman, FTIR and ^1^H-NMR) provided the information to characterize the synthetized biopolymer as PHB.

These findings underscore the capacity of a phototrophic microbiome, enriched with cyanobacteria, to achieve stable and long-term PHB production. Importantly, our research challenges traditional approaches relying on pure cultures by offering valuable insights into the application of phototrophic microbiomes and opens new frontiers in the field of sustainable PHB production. The implications of this work extend beyond the laboratory, paving the way for innovative solutions in meeting the growing demand for eco-friendly biopolymers.

## CRediT authorship contribution statement

**Beatriz Altamira-Algarra:** Conceptualization, Investigation, Writing – original draft. **Artai Lage:** Investigation. **Ana Lucía Meléndez:** Investigation. **Marc Arnau:** Investigation, Writing – original draft. **Eva Gonzalez-Flo:** Conceptualization, Supervision, Writing - review & editing. **Joan García:** Conceptualization, Supervision, Project administration, Funding acquisition, Writing - review & editing.

## Acknowledgements

This research was supported by the European Union’s Horizon 2020 research and innovation programme under the grant agreement No 101000733 (project PROMICON). B.A.A. thanks the Agency for Management of University and Research (AGAUR) for her grant [FIAGAUR_2021]. E.G.F. would like to thank the European Union-NextGenerationEU, Ministry of Universities and Recovery, Transformation and Resilience Plan for her research grant [2021UPF-MS-12]. J.G. acknowledges the support provided by the ICREA Academia program.

